# A 16S rRNA gene sequencing and analysis protocol for the Illumina MiniSeq platform

**DOI:** 10.1101/227108

**Authors:** Monica Pichler, Ömer K. Coskun, Ana Sofia Ortega, Nicola Conci, Gert Wörheide, Sergio Vargas, William D. Orsi

## Abstract

High-throughput sequencing of the 16S rRNA gene is widely used in microbial ecology, with Illumina platforms being widely used in recent studies. The MiniSeq, Illumina’s latest benchtop sequencer, enables more cost-efficient DNA sequencing relative to larger sequencing platforms *(e.g.* MiSeq). Here we used a modified custom primer sequencing approach to test the fidelity of the MiniSeq for high-throughput sequencing of the V4 hypervariable region of 16S rRNA genes from complex communities in environmental samples. To this end, we designed an additional sequencing primer that enabled application of a dual-index barcoding method on the MiniSeq. A mock community was sequenced alongside the environmental samples as a quality control benchmark. After careful filtering procedures, we were able to recapture a realistic richness of the mock community, and identify meaningful differences in alpha and beta diversity in the environmental samples. These results show that the MiniSeq can produce similar quantities of high quality V4 reads compared to the MiSeq, yet is a cost-effective option for any laboratory interested in performing high-throughput 16S rRNA gene sequencing.

**IMPORTANCE:** We modified a custom sequencing approach and used a mock community to test the fidelity of high-throughput sequencing on the Illumina MiniSeq platform. Our results show that the MiniSeq can produce similar quantities of high quality V4 reads compared to the MiSeq. In addition, our protocol increases feasibility for small laboratories to perform their own high-throughput sequencing of the 16S rRNA marker gene.

## Introduction

Continued improvements in DNA sequencing technologies have greatly helped in the democratization of sequencing (Tringe and Hugenholtz 2008) and high-throughput sequencing of the 16S rRNA marker gene is widely used to assess diversity and composition of microbial communities (Sogin et al. 2006; Huber et al. 2007; Bartram et al. 2011; Caporaso et al. 2012). However, the start-up and maintenance costs associated with high-throughput sequencing still hamper access to these technologies by smaller laboratories, many of which rely on sequencing centers and molecular core facilities to outsource high-throughput 16S rRNA gene sequencing.

Illumina’s MiniSeq benchtop platform enables cost-efficient high-throughput DNA sequencing relative to larger sequencing platforms *(e.g.* MiSeq). Thus, the goal of this study is to assess the quality of the MiniSeq generated data and to evaluate if the benchtop sequencer is a reliable and affordable option for any lab interested in performing 16S rRNA gene high-throughput sequencing. The acquisition cost for the MiniSeq starts at 50.000 € and yearly maintenance fees add up to approximately 5.000 €. Further, the 300 cycle Mid Output kit (for generating 2 x 150 bp paired-end reads) available for the MiniSeq is capable of generating up to 8 million pairs of reads, and the High Output version of this kit produces a volume of sequence data up to 25 million reads.

However, custom primer 16S rRNA sequencing protocols *(e.g.* Kozich et al. in 2013) were not designed for the MiniSeq and need to be adapted for this platform, in order to test their fidelity for 16S rRNA gene sequencing. Here, we modify an existing high-throughput 16S rRNA sequencing protocol using custom sequencing primers on the MiSeq (Kozich et al. 2013) to adapt this method for the new Illumina MiniSeq platform. We performed multiple high-throughput sequencing runs targeting the V4 hypervariable region of the 16S rRNA gene derived from complex environmental samples. Platform fidelity was assessed by alpha diversity analyses of a mock community of known species composition, which shows that with the proper quality controls the MiniSeq is capable of producing quality 16S rRNA gene sequence data at a reduced cost.

## Results and Discussion

In our study, we tested the fidelity of the Illumina MiniSeq platform for high-throughput sequencing of the 16S rRNA gene. We modified a paired-end sequencing strategy, which allowed for full length coverage of the hypervariable V4 region of the 16S rRNA gene. Because the V4 hypervariable region is ca. 250 bp in length, the 150 bp pair of reads produced by the MiniSeq overlap 50 bp on average.

The main modification of our MiniSeq protocol from the dual-index sequencing method of Kozich et al. 2013 is the use of an additional index sequencing primer. This additional index sequencing primer is necessary because the MiniSeq does not sequence the second index using adapters present on the flow cell surface as the MiSeq does. Rather, the MiniSeq reads Index 2 only after the clusters have been turned around to sequence the paired-end reads (Figure 1). Thus, in addition to the three sequencing primers described by Kozich et al. (2013), we designed and used an Index 2 sequencing primer (5’-TTACCGCGGCKCGTGGCACACAATTACCATA-3’) (see Table 1) to enable the dual-index barcoding method on the MiniSeq. We tested this modified approach on three different 16S rRNA sequencing runs including diverse environmental samples as well as a mock community composed of 18 different bacterial species. The mock community was created from pure cultures, whose 16S rRNA genes were determined through Sanger sequencing to be >3% different (Table S1). Environmental samples were collected from salt marsh sediments, freshwater pond sediments, marine sponges, and salt water aquaria.

**Figure 1.**
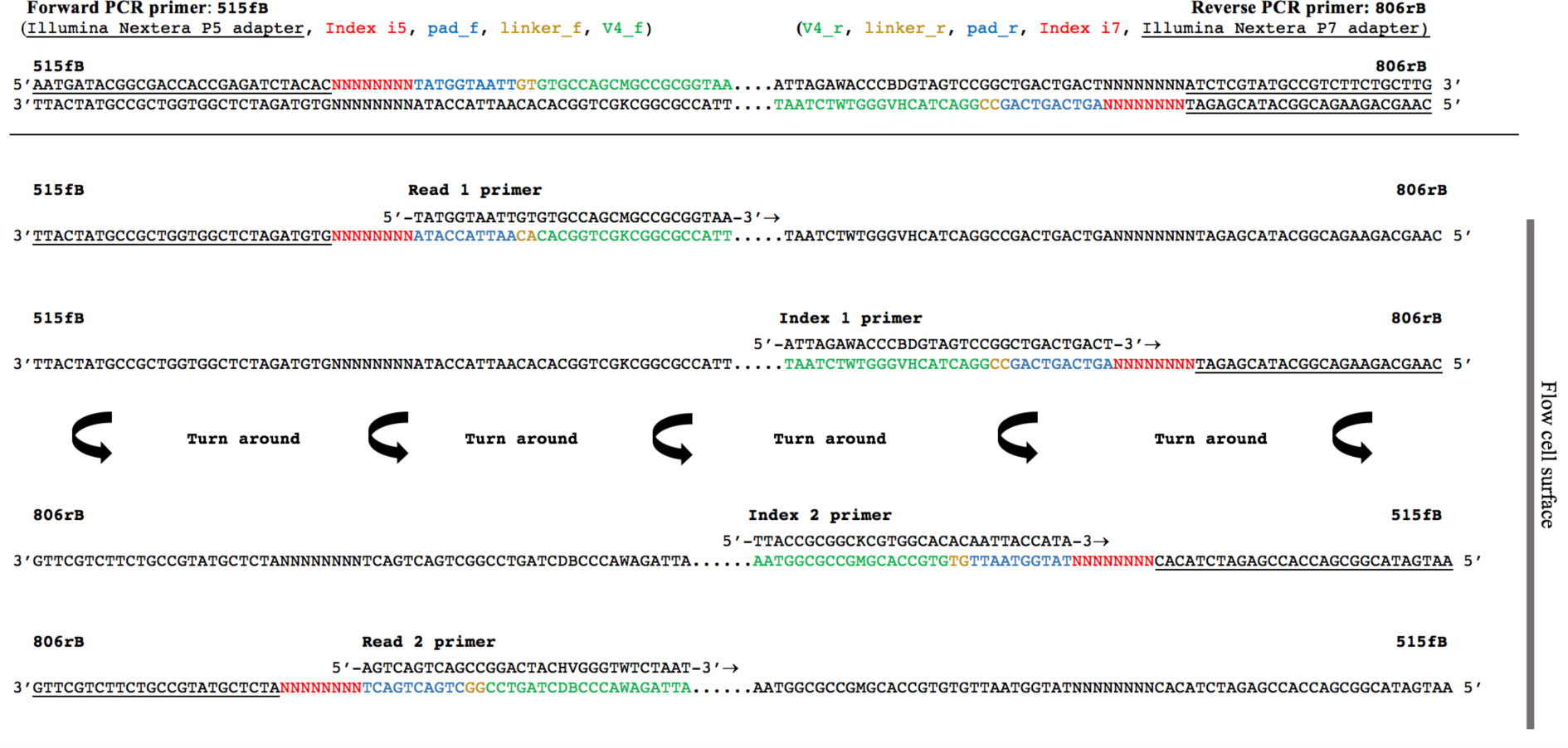
Schematic description of the dual-index sequencing strategy on the MiniSeq. Reading the figure from top to bottom shows the sequential order of paired-end sequencing steps (four total). “Turn around” indicates the step of paired-end turn around on the flow cell surface. The sequencing proceeds in the direction of the flow cell surface, which in this figure is located on the right side (arrows point in direction of sequencing reaction). Sequencing starts by using Read 1 primer to sequence Read 1, followed by Index 1 primer to generate Index 1. The MiniSeq only uses the oligonucleotides on the flow cell for bridging and both the second index and the paired read are sequenced after the clusters are turned around. Hence an Index 2 primer is needed to sequence Index 2. Read 2 is then sequenced by using the Read 2 primer (after Kozich et al. 2013).

**Table 1.**
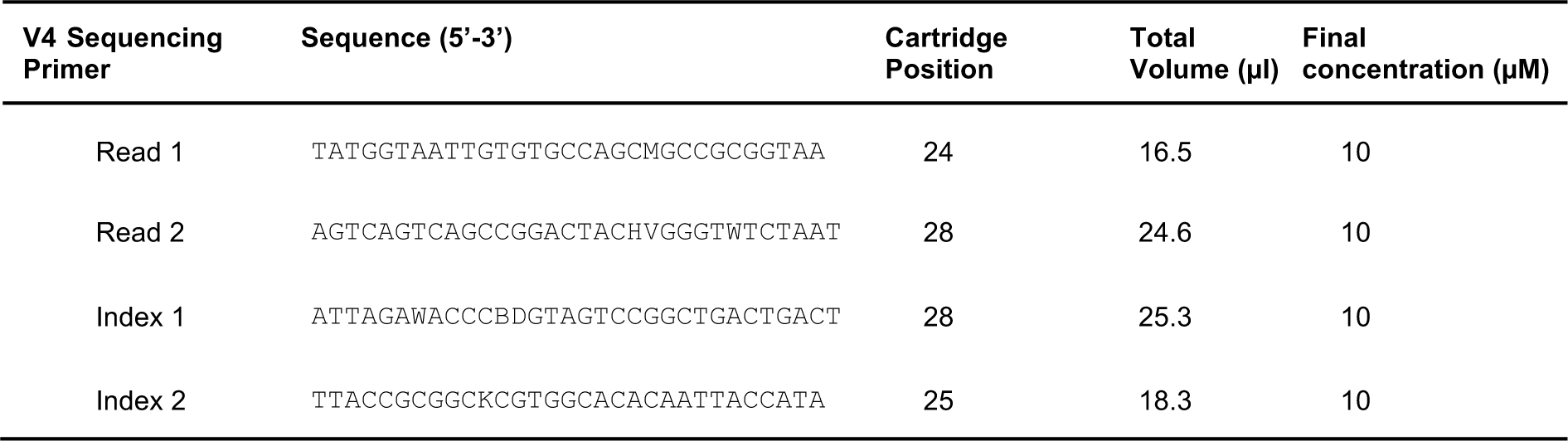
Custom sequencing primers used to target the V4 region. The primers were diluted and loaded into the correct cartridge position.

### Run performances

The MiniSeq performed 151 cycles of both forward and reverse reads (see Table 2 for all run metrics). Run A yielded a total of 1.23 Gbp with cluster density of 76 ± 9 K/mm^2^ detected by image analysis and 73.28 ± 13.91% of the clusters passing filter (PF) on the platform. Hence, 5 million clusters were generated, of which approximately 3.9 million passed the filter. 92.10% of all bases from both reads were assigned a quality score of Q ≥ 30. About 8% of all reads were aligned to the quality control PhiX genome and removed. The calculated sequencing error rate for the MiniSeq can be calculated as the percentage of PhiX reads (from the spiked in sample) with mismatches to the PhiX genome. This was a preliminary indication that the MiniSeq had an error rate of 1.37%. Sequencing run B generated 3.31 Gbp and achieved optimal cluster density of 170 ± 3 K/mm^2^ with 85.65 ± 1.28% of clusters PF. In total, 12.2 million clusters were generated, of which 10.5 million passed filtering. 88.61% of all bases from both reads were assigned a quality score of Q ≥ 30. About 25% of the cluster passing filtering could be aligned to the PhiX genome, which resulted in a calculated error rate of 0.79%. Run C yielded 2.67 Gbp with a cluster density of 124 ± 1K/mm^2^ and 95.52 ± 0.54% of the clusters PF (8.5 million of 8.9 million clusters). A quality score of Q ≥ 30 was achieved by 94.79% of all bases. 12% of all reads were aligned to PhiX and an error rate of 0.43 was indicated.

Sequencing run A appeared to be under-clustered considering the low cluster density. According to Illumina’s specifications (Illumina 2016), the recommended cluster density for the mid-output kit (300 cycles) on the MiniSeq is 170-220 K/mm^2^ (slightly below for low diversity libraries). Hence, we optimized cluster density by increasing the genetic diversity of the samples for sequencing runs B and C, by spiking in an additional Illumina library of genomic DNA from a marine sponge (see Methods). This further resulted in clusters PF>80% expected for optimized cluster density on the platform.

**Table 2.**
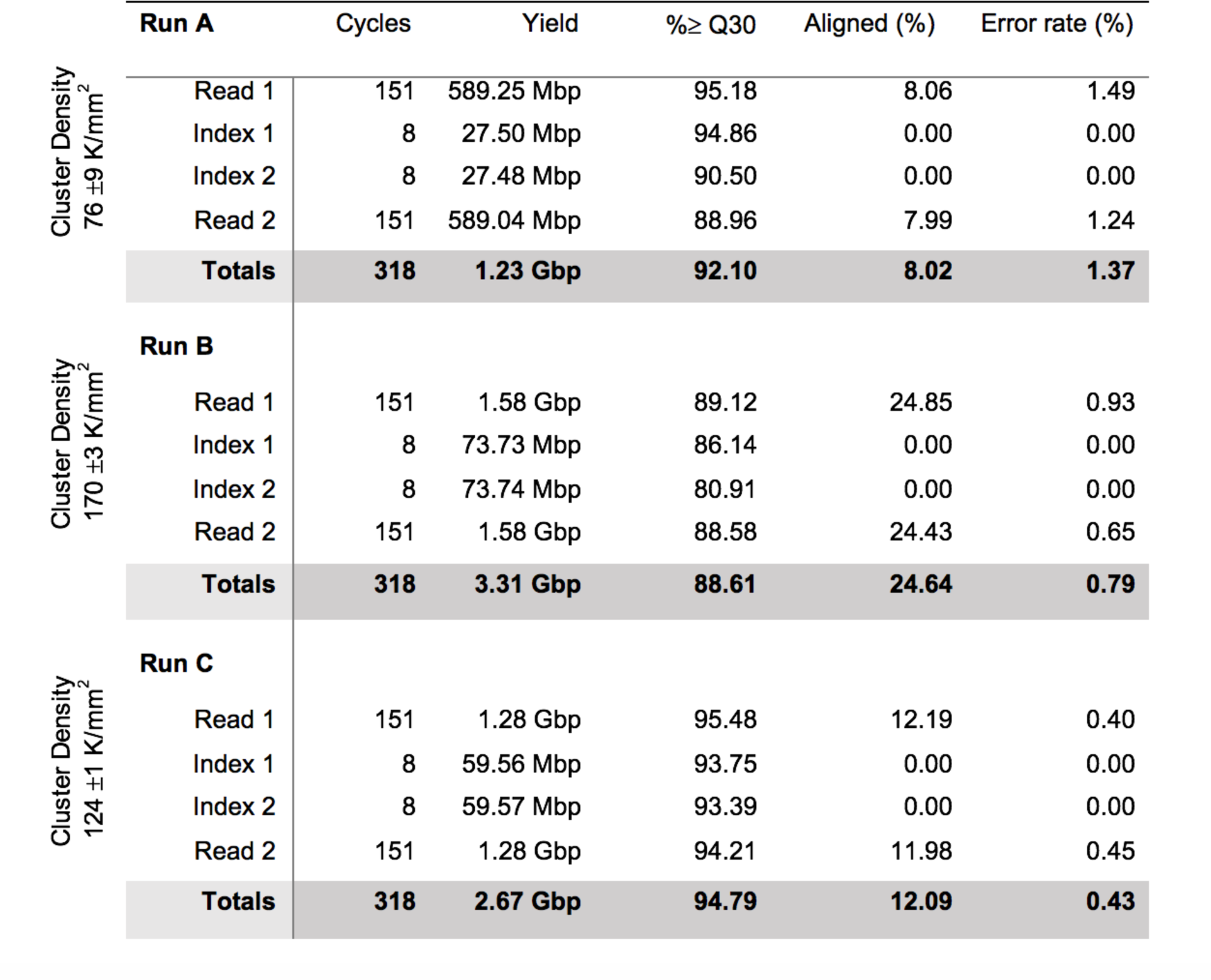
Overview of the 16S rRNA sequencing run metrics.

|  | Run A | Cycles | Yield | %≥ Q30 | Aligned (%) | Error rate (%) |
| --- | --- | --- | --- | --- | --- | --- |
| Cluster Density<br>76 ±9 K/mm<sup>2</sup> | Read 1 | 151 | 589.25 Mbp | 95.18 | 8.06 | 1.49 |
|  | Index 1 | 8 | 27.50 Mbp | 94.86 | 0.00 | 0.00 |
|  | Index 2 | 8 | 27.48 Mbp | 90.50 | 0.00 | 0.00 |
|  | Read 2 | 151 | 589.04 Mbp | 88.96 | 7.99 | 1.24 |
|  | Totals | 318 | 1.23 Gbp | 92.10 | 8.02 | 1.37 |
| Cluster Density<br>170 ±3 K/mm<sup>2</sup> | Run B |  |  |  |  |  |
|  | Read 1 | 151 | 1.58 Gbp | 89.12 | 24.85 | 0.93 |
|  | Index 1 | 8 | 73.73 Mbp | 86.14 | 0.00 | 0.00 |
|  | Index 2 | 8 | 73.74 Mbp | 80.91 | 0.00 | 0.00 |
|  | Read 2 | 151 | 1.58 Gbp | 88.58 | 24.43 | 0.65 |
|  | Totals | 318 | 3.31 Gbp | 88.61 | 24.64 | 0.79 |
| Cluster Density<br>124 ±1 K/mm<sup>2</sup> | Run C |  |  |  |  |  |
|  | Read 1 | 151 | 1.28 Gbp | 95.48 | 12.19 | 0.40 |
|  | Index 1 | 8 | 59.56 Mbp | 93.75 | 0.00 | 0.00 |
|  | Index 2 | 8 | 59.57 Mbp | 93.39 | 0.00 | 0.00 |
|  | Read 2 | 151 | 1.28 Gbp | 94.21 | 11.98 | 0.45 |
|  | Totals | 318 | 2.67 Gbp | 94.79 | 12.09 | 0.43 |

### Terminal G homopolymers

The MiniSeq uses a 2-channel sequencing by synthesis (SBS) method compared to the 4-channel SBS technology used on the MiSeq and HiSeq instruments. Clusters appearing in red and green are cytosine (C) and thymine (T) nucleotides, respectively, while adenine (A) bases are detected in both channels and appear yellow. Guanine (G) nucleotides are unlabelled clusters and are seen in neither channel hence they appear black (Illumina a).

In our first 16S rRNA sequencing run (run A), 7% of forward reads and 8% of reverse reads had long (>10) terminal poly-G strings (see Figure S1). As G indicates lack of sequencing signal with the Illumina 2-dye chemistry *(e.g.* black), this may be due to underclustering on the flow cell, low diversity in the 16S libraries, or partially amplified V4 PCR fragments carried over during the gel extraction. This phenomenon appears to be due to the low diversity inherent in 16S sequencing datasets, as this was not observed in any of our prior genome or transcriptome sequencing libraries on the MiniSeq (data not shown). Long poly-G strings were also not detected in the data from the other 16S sequencing runs (run B and C), which had genomic DNA spiked in to increase the nucleotide diversity. Thus, we recommend that researchers mix separately indexed genomic libraries together with their 16S rRNA gene libraries when sequencing on the MiniSeq to reduce the number of terminal G homopolymers.

We removed all sequences with G homopolymers >10 nucleotides prior to data analysis. As an additional precaution, we removed all OTUs that were represented by <10 sequences, which may have contained spurious guanine homopolymers shorter than <10 nucleotides. We urge caution when analyzing rare taxa (Sogin et al. 2006) with 16S data generated on the MiniSeq, as sequences with terminal poly-G homopolymers need to be carefully accounted for. In order to determine whether any remaining terminal poly-G homopolymers not removed by the above quality controls *(e.g.* those less than 10 residues) affected the true 16S diversity, we compared the number of OTUs in the mock community to the true richness.

### OTU assignments

In order to test the fidelity of the MiniSeq for 16S rRNA gene sequencing, we clustered OTUs from the mock community dataset using the USEARCH pipeline (Edgar 2010). After these data processing steps, and removal of chimeric sequences (see Methods), the UPARSE algorithm (Edgar 2013) recovered 17 out of the 18 species in our mock community and 4 spurious OTUs in run A, and 15 out of 18 plus 4 spurious in run C (Figure 2). While the 16S mock community was not sequenced alongside the environmental samples in run B, it was used to analyse the generated data set. Again, the number of species found in the mock community was close to its true composition (16 out of 18 species, 4 spurious OTUs). Thus, the UPARSE method could accurately recover the microbial richness from our MiniSeq 16S rRNA gene data. Other studies *(e.g.* Edgar 2013; Flynn et al. 2015) also showed that the number of OTUs generated with UPARSE is in close concordance with the number of species in a mock community. While the exact number of OTUs in the mock community was not obtained with UPARSE, mock communities are rarely recovered at the exact richness after 16S high-throughput sequencing with variability reaching >30% of the richness in the original mock community even under stringent criteria (Edgar 2013). This is typically attributed to additional undetected contaminants, and single sequencing errors that can occur in low abundance in the sample index barcodes (Edgar 2013). Our quality control procedures for the MiniSeq 16S rRNA gene data appears to be sufficiently prudent, because the richness of our recovered mock community OTUs relative to the starting richness falls within the variability of stringently controlled mock community sequence analyses (Edgar 2013). To control for contamination, we also sequenced lab dust samples and extraction blanks and removed OTUs shared with the environmental samples. After removal of contaminant OTUs, a significantly different (ANOSIM: P=0.001, R: 0.8) microbiome for each sample was observed (Figure 3). Given that the richness of the mock community is close to the true value, these beta diversity analyses show that the MiniSeq is a viable platform for high-throughput 16S rRNA gene sequencing studies of microbiomes.

**Figure 2.**
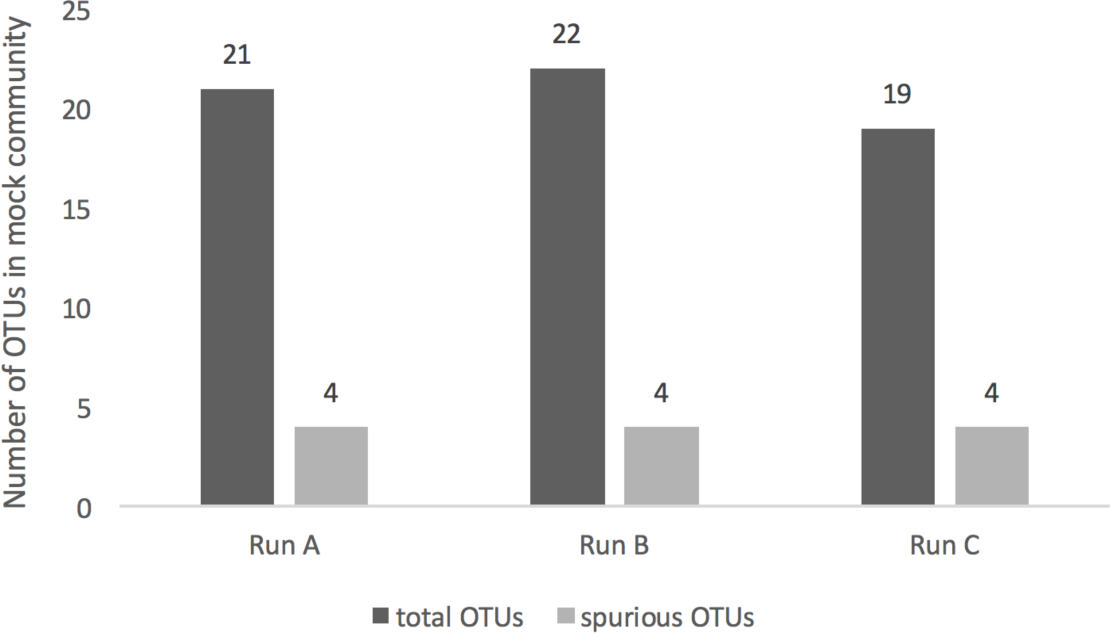
OTU assessment for the mock community composed of 18 defined species. UPARSE generated an accurate estimate of the microbial community in all performed 16S rRNA sequencing runs, given the low number of spurious OTUs.

**Figure 3.**
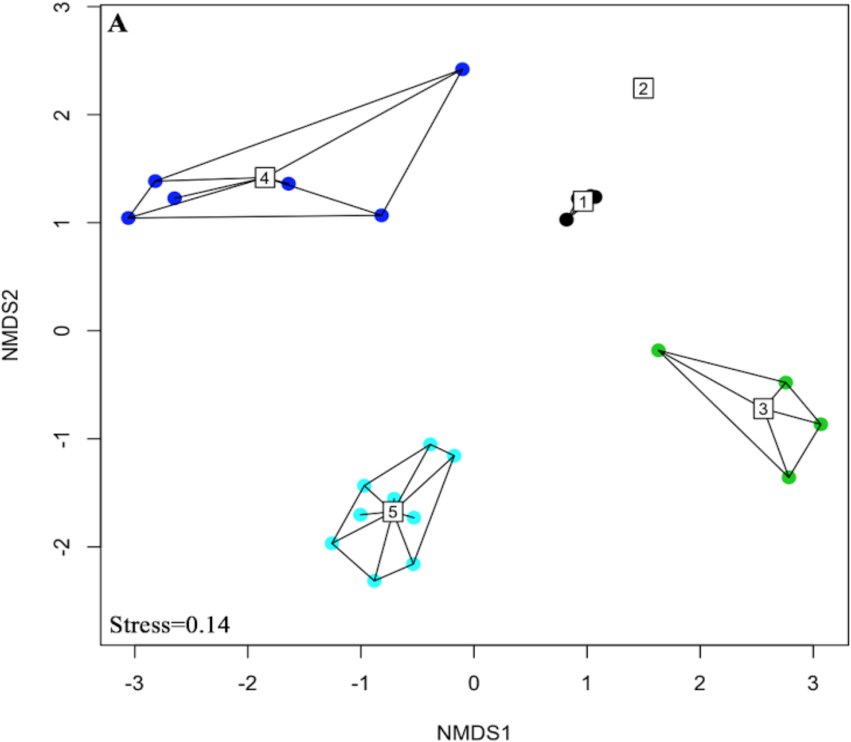
Non-metric multidimensional scaling analysis showing microbial beta diversity of the 16S data sets. (1) mock community replicates, (2) salt water aquaria, (3) marine sponge, (4) pond sediments, (5) salt marsh sediments.

Comparing sequencing fidelity across platforms is a feasible way of validating high-throughput sequencing approaches (Caporaso et al. 2012). However, mock communities can also be used as a way to test the fidelity of high-throughput sequencing platforms (Benítez-Páez, Portune, and Sanz 2016; Caporaso et al. 2011). Thus, while we do not compare our results to those obtained from larger sequencing platforms *e.g.* a MiSeq (as described by Caporaso et al. 2012), the analyses of the mock community show that the MiniSeq is able to capture a realistic picture of its microbial diversity. With our results, we evaluated the MiniSeq as a reliable and affordable alternative to larger sequencing platforms. Our protocol thus increases feasibility for small laboratories to perform their own high-throughput sequencing of the 16S rRNA marker gene.

## Material and Methods

### Cultivation and DNA Extraction of the 16S mock community

To create a mock community (>3% dissimilarity threshold, see Table S1), pure cultures were isolated from soil, human skin, cell phone swabs, freshwater and saltwater, and grown on agar plates for 3-7 days at room temperature. For genomic DNA extraction, a small amount of each bacterial strain was transferred into a 2 mL sterile lysing Matrix E tube and 800 μl of preheated (60°C) sterile filtered C1 extraction buffer (38 mL saturated NaPO4 [1M] buffer, 7.5 mL 100% ethanol, 4 mL MoBio’s lysis buffer solution C1 [MoBio, Carlsbad, CA], 0.5 mL 10% SDS) was added. The samples were homogenized for 40 sec at a speed of 6 m/sec using a QuickPrep-24 5G homogenizer (MP Biomedicals, Santa Ana, CA) and heated for 2 min at 99°C in an Eppendorf ThermoMixer C (Thermo Fisher Scientific, Waltham, MA), followed by two freeze-thaw (-80°C/room temperature) cycles to lyse bacterial cells. After repetition of the homogenizing step, the samples were centrifuged for 10 min at 14.800 rpm in a Heraeus Pico 21 centrifuge (Thermo Fisher Scientific, Waltham, MA). Microbial DNA was purified using the MoBio PowerClean Pro DNA Clean-Up Kit (Qiagen, Hilden, Germany) following the manufacturer’s instructions using 100 μl of the supernatant. DNA was quantified fluorometrically on the Qubit version 3.0 (Life Technologies, Grand Island, NY) using the Qubit dsDNA high sensitivity assay kit (Life Technologies).

To confirm the number of species in the mock community, the full length 16S rRNA gene of each isolate was amplified and sequenced by Sanger sequencing. Two conserved primers (27f, 1492r) were used to amplify the entire gene during PCR with the following conditions: initial denaturation at 95°C for 3 min; 30 cycles of denaturation at 95°C for 30 sec; annealing at 56°C for 30 sec; elongation at 72°C for 1 min and a final 5 min extension at 72°C. Individual reactions consisted of 1 μl template DNA, 5 μl 5x Green GoTaq Flexi Buffer (Promega), 3 μl MgCl_2_ (25mM), 1 μl fw primer (10 μM), 1 μl rv primer (10 μM), 12.9 μl nuclease-free water, dNTP Mix (10mM) and 0.1 μl GoTaq Green DNA Polymerase (Promega). The amplicons were subjected to Sanger sequencing using the facilities of the Biocenter of the Ludwig-Maximilian University (LMU), Martinsried. To confirm dissimilarity thresholds of >3% for all 18 species, we aligned the sequences using BLAST (Altschul et al. 1990). We pooled the isolates at equimolar concentration and created technical replicates of the mock community to assess the reproducibility of the method.

Genomic DNA of contaminants, comprising of dust samples (n=9) and extractions blanks (n=3), and all other environmental samples (run A, n=30; run B, n=88; run C, n=83) was extracted following the same method, but with an additional step. Before purification, the supernatant was concentrated to approximately 100 μl in 50 MW KDa Amicon filters by centrifuging for 15 min at 47000 rpm using the Allegra X-30R centrifuge (Beckman Coulter, Brea, CA) to improve DNA yield. The contaminants were collected from three different laboratory rooms of the LMU building. Environmental samples included salt marsh sediments, freshwater pond sediments, marine sponges, and salt water aquaria.

### 16S amplicon library preparation

We followed the dual-index paired-end sequencing approach previously described by Kozich et al. (2013) and developed for sequencing on the Illumina MiSeq platform. The V4 region of the 16S rRNA gene was amplified with unique barcoded PCR primers 515fB (5’ - AAT GAT ACGGCGACCACCGAGAT CT ACAC NNNNNNNN **TATGGTAATT** GT *GTGCCAGCMGCCGCGGTAA* - 3’) and 806rB (5’ - CAAGCAGAAGACGGCATACGAGAT NNNNNNNN **AGTCAGTCAG** CC *GGACTACHVGGGTWTCTAAT* - 3’) (see Table S2 in the supplemental material for barcodes). For the third 16S rRNA sequencing run (run C), we used modified 515f/806rB primer constructs (515f: GTGYCAGCMGCCGCGGTAA; 806rB: GGACTACNVGGGTWTCTAAT), which include the latest changes that increase coverage of Thaumarchaeota (Walters et al. 2015) and further enable capturing of a greater diversity of the marine SAR11 clade (Apprill et al. 2015). The primer sequences consist of the appropriate Illumina adapter (P5 or P7; underlined) complementary to the oligonucleotides on the flow cell, an 8-nt index sequence representing the unique barcode for every sample (N region), a 10-nt pad sequence (bold), a 2-nt linker (GT, CC) and the specific primer for the V4 region (italic) (Kozich et al. 2013). All samples were amplified on the Biometra TProfessional Thermocycler (Biometra, Göttingen, Germany) in a total reaction volume of 24 μl including 2 μl template DNA, 5 μl 5x Green GoTaq Flexi Buffer (Promega), 1 μl forward primer (10 μM), 1 μl reverse primer (10 μM), 1μl dNTP Mix (10mM), 3 μl MgCl_2_ (25mM), 0.2 μl GoTaq Green DNA Polymerase (Promega) and 12.8 μl nuclease-free water. PCR program was run as follows: initial denaturation at 95°C for 3 min, followed by 30 cycles of denaturation at 95°C for 30 sec, annealing at 56°C for 30 sec, elongation at 72°C for 1 min and a final elongation step at 72°C for 5 min.

The barcoded DNA amplicons were analysed on a 1.5% (w/v) agarose gel, and excised and purified for sequencing using the Zymoclean Gel DNA Recovery Kit (Zymo Research, Irvine, CA), adding 15 μl of buffer EB to elute DNA. After gel extraction, DNA concentrations were measured using Qubit and diluted first to 10 nM and then to a final 1 nM in a serial dilution before the samples were pooled (adding 5 μl of every sample).

### 16S sequencing strategy and primer design

We performed three paired-end 16S rRNA sequencing runs on the MiniSeq (run A, B and C). For all runs, we used the MiniSeq Mid Output Reagent Kit (300 cycles) including a reagent cartridge, a single-use flow cell and hybridization buffer HT1. To prepare our normalized amplicon libraries for sequencing, we followed the MiniSeq Denature and Dilute Libraries Guide (Protocol A) (Illumina b) with some customizations. For run A, we combined 500 μl of the denatured and diluted 16S library (1.8 pM) with 20 μl of denatured and diluted Illumina generated PhiX control library (1.8 pM) to increase the diversity of the low nucleotide pool and to assess sequencing error rates.

For run B and C, we combined 350 μl of the 16S library (1.8 pM) with 150 μl of a denatured and diluted genomic sponge library *(Ephydatia fluviatilis,* 1.8 pM) and additionally added 15 μl of PhiX (1.8 pM). The final 1.8 pM libraries were loaded into the “Load samples” well of the reagent cartridge. For each run, we used four custom sequencing primers Read 1,Index 1, Index 2 and Read 2, which were diluted and loaded into the correct position of the reagent cartridge (see Table 1).

We had to design an additional Index 2 sequencing primer (see Table 1) to enable the dual-index barcoding method on the MiniSeq. This additional index sequencing primer is needed because, as opposed to the MiSeq, the MiniSeq only reads Index 2 after the clusters have been turned around to sequence the pair reads (see Figure 1). Sequencing proceeds in the direction of the flow cell and starts by generating Read 1 (150 bp) using Read 1 sequencing primer, followed by obtaining Index 1 (8 bp) using Index 1 sequencing primer. Clusters were turned around by using the oligonucleotides provided on the flow cell. After bridging, Index 2 sequencing primer generates Index 2 (8 bp) and Read 2 sequencing primer finally obtains Read 2 (150 bp).

### 16S bioinformatics analyses and OTU assignment

Demultiplexing and base calling were both performed using bcl2fastq Conversion Software v2.18 (Illumina, Inc.). All bioinformatics analysis were conducted in USEARCH version 9.2.64 (Edgar 2010) and QIIME version 1.9.1 (Caporaso et al. 2010). The initial step was to assemble paired-end reads using the fastq_merge pairs command with default parameters allowing for a maximum of five mismatches in the overlapping region. Stringent quality filtering was carried out using the fastq_filter command. We discarded low quality reads by setting the maximum expected error threshold (E_max), which is the sum of the error probability provided by the Q score for each base, to 1. Reads were de-replicated and singletons discarded. Reads were clustered into OTUs sharing 97% sequence identity using the heuristic clustering algorithm UPARSE (Edgar 2013), which is implemented in the cluster_otus command. The algorithm performs *de novo* chimera filtering and OTU clustering simultaneously (Edgar 2013). The usearch_global command assigned the reads to OTUs and created an OTU table for further downstream analysis. Taxonomy was assigned in QIIME through BLASTn searches (Altschul et al. 1990) against the SILVA ribosomal RNA gene database (Quast et al. 2013). The OTU table was rarefied in QIIME to the sample with the least number of reads using the single_rarefaction.py command. This required a conversion of the OTU table text file into biom (biological observation matrix) format using the convert biom command. As a quality control step, we removed all OTUs containing <10 sequences and which had no BLASTn hit.

### 16S data analysis

In order to investigate beta diversity structures of our samples, we performed downstream analysis in R version 3.3.0 (R Development Core Team 2011). Non-metric multivariate (NMDS) analyses of the microbial communities were calculated using a Bray Curtis distance in the Vegan package (Oksanen et al. 2017). Analysis of Similarity (ANOSIM) was performed using 999 permutations with a Bray Curtis distance.

## Acknowledgements

This manuscript is an outcome of a research project carried out by MP within the framework of the international Master’s program Geobiology & Paleobiology at the Faculty of Geosciences of the Ludwig-Maximilians-Universität München.

## Supplementary Material

**Table S1.** Composition of the artificially created 16S mock community. Taxa could not be determined to the exact species level, yet all isolates show < 97% similarity cut-off for species differentiation.

| No. | Bacterial Isolate (<97% similarity) |
| --- | --- |
| 1 | *Staphylococcus* sp. |
| 2 | *Bacillus* sp. |
| 3 | *Bacillus* sp. |
| 4 | *Micrococcus* sp. |
| 5 | *Acinetobacter* sp. |
| 6 | *Enterobacter* sp. |
| 7 | *Aeromonas* sp. |
| 8 | *Carnobacterium* sp. |
| 9 | *Exiguobacterium* sp. |
| 10 | *Janthinobacterium* sp. |
| 11 | *Pseudomonas* sp. |
| 12 | *Photobacterium* sp. |
| 13 | *Pseudoalteromonas* sp. |
| 14 | *Vibrio* sp. |
| 15 | *Rhodococcus* sp. |
| 16 | *Sphingobium* sp. |
| 17 | *Arthrobacter* sp. |
| 18 | *Mycobacterium* sp. |

**Table S2.** Barcoded primer combinations.

| Forward Primer | Barcode (i5) | Reverse Primer | Barcode (i7) |
| --- | --- | --- | --- |
| 515F.A501 | ATCGTACG | 806RB.A701 | AACTCTCG |
| 515F.A502 | ACTATCTG | 806RB.A702 | ACTATGTC |
| 515F.A503 | TAGCGAGT | 806RB.A703 | AGTAGCGT |
| 515F.A504 | CTGCGTGT | 806RB.A704 | CAGTGAGT |
| 515F.A505 | TCATCGAG | 806RB.A705 | CGTACTCA |
| 515F.A506 | CGTGAGTG | 806RB.A706 | CTACGCAG |
| 515F.A507 | GGATATCT | 806RB.A707 | GGAGACTA |
| 515F.A508 | GACACCGT | 806RB.A708 | GTCGCTCG |

**Figure S1.**
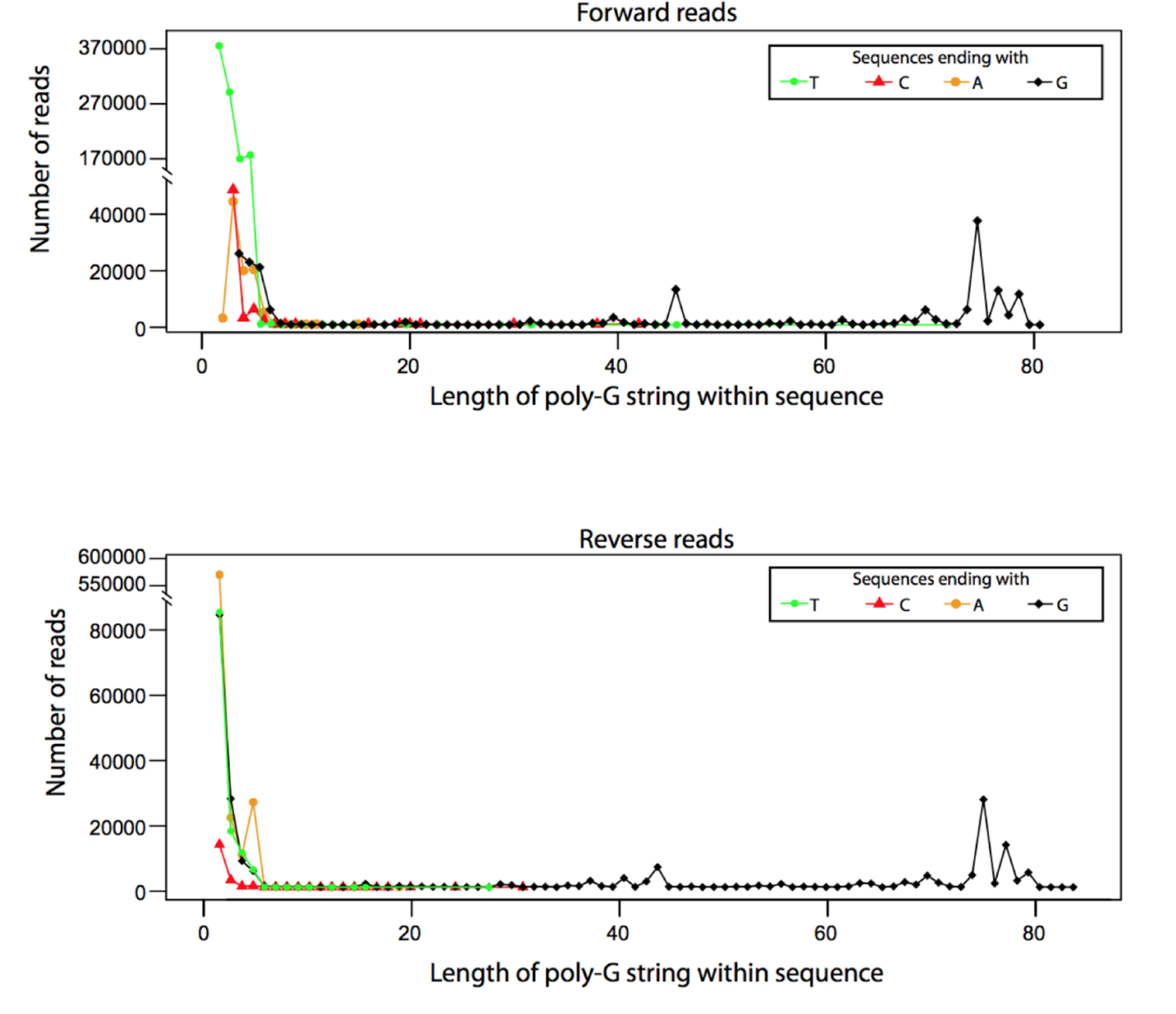
Plots showing the abundance of homopolymeric guanine repeats of different length in forward and reverse reads (Run A). Note that ca. 7% of reads exhibit long (>10 nucleotides) homopolymers of G’s, most of which tended to be between 75-80 nucleotides. These erroneous homopolymers appear to be mostly restricted to the ends of the sequence (not internal G homopolymers), as sequences ending with A, T, and C did not have long (>10) poly-G homopolymers. These homopolymers were not observed in genome and transcriptome datasets sequenced on the MiniSeq (data not shown), and are likely due to a combination of the Illumina 2-dye chemistry and the relatively low diversity of 16S libraries.

## References

Altschul, S. F., W. Gish, W. Miller, E. W. Myers and D. J. Lipman. 1990. “Basic Local Alignment Search Tool.” Journal of Molecular Biology 215 (3): 403–10.

Apprill, A., S. McNally, R. Parsons, and L. Weber. 2015. “Minor Revision to V4 Region SSU rRNA 806R Gene Primer Greatly Increases Detection of SAR11 Bacterioplankton.” Aquatic Microbial Ecology: International Journal 75 (2): 129–37.

Bartram, Andrea K., Michael D. J. Lynch, Jennifer C. Stearns, Gabriel Moreno-Hagelsieb, and Josh D. Neufeld. 2011. “Generation of Multimillion-Sequence 16S rRNA Gene Libraries from Complex Microbial Communities by Assembling Paired-End Illumina Reads.” Applied and Environmental Microbiology 77 (11): 3846–52.

Benítez-Páez, Alfonso, Kevin J. Portune, and Yolanda Sanz. 2016. “Species-Level Resolution of 16S rRNA Gene Amplicons Sequenced through the MinION™ Portable Nanopore Sequencer.” GigaScience 5 (January): 4.

Caporaso, J. Gregory, Justin Kuczynski, Jesse Stombaugh, Kyle Bittinger, Frederic D. Bushman, Elizabeth K. Costello, Noah Fierer, et al. 2010. “QIIME Allows Analysis of High-Throughput Community Sequencing Data.” Nature Methods 7 (5). Nature Publishing Group: 335–36.

Caporaso, J. Gregory, Christian L. Lauber, William A. Walters, Donna Berg-Lyons, James Huntley, Noah Fierer, Sarah M. Owens, et al. 2012. “Ultra-High-Throughput Microbial Community Analysis on the Illumina HiSeq and MiSeq Platforms.” The ISME Journal 6 (8). Nature Publishing Group: 1621–24.

Caporaso, J. Gregory, Christian L. Lauber, William A. Walters, Donna Berg-Lyons, Catherine A. Lozupone, Peter J. Turnbaugh, Noah Fierer, and Rob Knight. 2011. “Global Patterns of 16S rRNA Diversity at a Depth of Millions of Sequences per Sample.” Proceedings of the National Academy of Sciences of the United States of America 108 Suppl 1 (March): 4516–22.

Edgar, Robert C. 2010. “Search and Clustering Orders of Magnitude Faster than BLAST.” Bioinformatics 26 (19): 2460–61.

Edgar, Robert C. 2013. “UPARSE: Highly Accurate OTU Sequences from Microbial Amplicon Reads.” Nature Methods 10 (10): 996–98.

Huber, Julie A., David B. Mark Welch, Hilary G. Morrison, Susan M. Huse, Phillip R. Neal, David A. Butterfield, and Mitchell L. Sogin. 2007. “Microbial Population Structures in the Deep Marine Biosphere.” Science 318 (5847): 97–100.

Illumina, 2016. Optimizing Cluster Density on Illumina Sequencing Systems, Available at: https://support.illumina.com/content/dam/illumina-marketing/documents/products/other/miseq-overclustering-primer-770-2014-038.pdf.

Kozich, James J., Sarah L. Westcott, Nielson T. Baxter, Sarah K. Highlander, and Patrick D. Schloss. 2013. “Development of a Dual-Index Sequencing Strategy and Curation Pipeline for Analyzing Amplicon Sequence Data on the MiSeq Illumina Sequencing Platform.” Applied and Environmental Microbiology 79 (17): 5112–20.

Oksanen, J. F., G. Blanchet, M. Friendly, R. Kindt, P. Legendre, D. McGlinn, P. R. Minchin, et al. 2017. Vegan: Community Ecology Package. R Package Version 2.4-2 (version version 2.4-2). https://CRAN.R-project.org/package=vegan.

Quast, Christian, Elmar Pruesse, Pelin Yilmaz, Jan Gerken, Timmy Schweer, Pablo Yarza, Jörg Peplies, and Frank Oliver Glöckner. 2013. “The SILVA Ribosomal RNA Gene Database Project: Improved Data Processing and Web-Based Tools.” Nucleic Acids Research 41 (Database issue): D590–96.

R Development Core Team. 2011. R: The R Project for Statistical Computing. https://www.r-project.org/.

Sogin, Mitchell L., Hilary G. Morrison, Julie A. Huber, David Mark Welch, Susan M. Huse, Phillip R. Neal, Jesus M. Arrieta, and Gerhard J. Herndl. 2006. “Microbial Diversity in the Deep Sea and the Underexplored ‘Rare Biosphere.’” PNAS 103 (32): 12115–20.

Tringe, Susannah G., and Philip Hugenholtz. 2008. “A Renaissance for the Pioneering 16S rRNA Gene.” Current Opinion in Microbiology 11 (5): 442–46.

Walters, William, Embriette R. Hyde, Donna Berg-Lyons, Gail Ackermann, Greg Humphrey, Alma Parada, Jack A. Gilbert, et al. 2015. “Improved Bacterial 16S rRNA Gene (V4 and V4-5) and Fungal Internal Transcribed Spacer Marker Gene Primers for Microbial Community Surveys.” mSystems 1 (1). doi:10.1128/mSystems.00009-15.

